# Whole-head recording of chemosensory activity in the marine annelid *Platynereis dumerilii*

**DOI:** 10.1101/391920

**Authors:** Thomas Chartier, Joran Deschamps, Wiebke Dürichen, Gáspár Jékely, Detlev Arendt

## Abstract

Chemical detection is key to various behaviours in both marine and terrestrial animals. Marine species, though highly diverse, have been underrepresented so far in studies on chemosensory systems, and our knowledge mostly concerns the detection of airborne cues. A broader comparative approach is therefore desirable. Marine annelid worms with their rich behavioural repertoire represent attractive models for chemosensory studies. Here, we study the marine worm *Platynereis dumerilii* to provide the first comprehensive study of head chemosensory organ physiology in an annelid. By combining microfluidics and calcium imaging, we record neuronal activity in the entire head of early juveniles upon chemical stimulation. We find that *Platynereis* uses four types of organs to detect stimuli such as alcohols, esters, amino acids and sugars. Antennae, but not nuchal organs or palps as generally hypothesised in annelids, are the main chemosensory organs. We report chemically-evoked activity in possible downstream brain regions including the mushroom bodies, which are anatomically and molecularly similar to insect mushroom bodies. We conclude that chemosensation is a major sensory modality for marine annelids, and propose early *Platynereis* juveniles as a model to study annelid chemosensory systems.

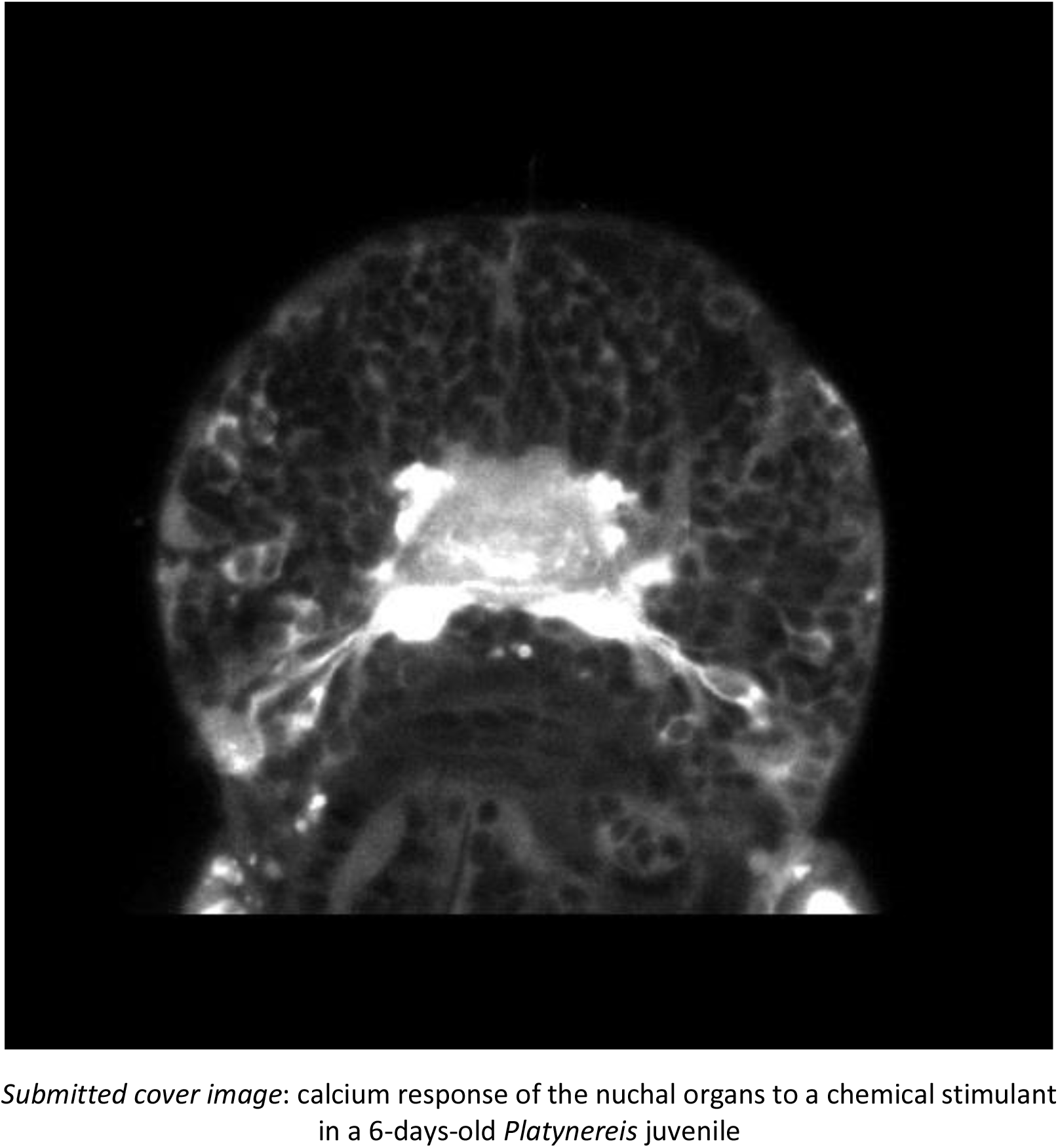

## Introduction

Chemical signals are central to animal behaviour, including feeding, predation, courtship and mating, aggregation, defence, habitat selection, communication [1]. Adapting to variable habitats and changing chemical landscapes, animals have evolved a broad variety of chemosensory organs. Investigations of chemosensory systems in mammals, insects and nematodes have provided insights into the molecular and cellular basis of how chemical information is encoded into neuronal activity [2–4]. While similar circuit architectures can be found in distant species at some steps of information processing, this appears to be no general rule [5, 6]. Genomic studies have revealed that receptor proteins are highly diverse in the animal kingdom [7], and can be entirely different between distant species - vertebrates and insects for example use distinct types of receptors [8, 9]. Hence, a broader comparative approach will facilitate the elucidation of both general operating principles and evolutionary origins of animal chemosensation. Despite studies in fish and crustaceans [10, 11], and to a lesser extent in molluscs [12, 13], our current understanding of animal chemosensation still mainly concerns terrestrial and airborne cues. Marine animals thus deserve more attention.

Marine annelids, traditionally referred to as ‘polychaetes’, represent an attractive group for chemosensory studies. These worms, represented by more than 10.000 species, are typically free-living, burrow in the marine sediment, or build tubes. They are known to respond to chemical signals in reproduction, feeding, aggression, avoidance, aggregation, environment probing, larval settlement and metamorphosis [14]. Marine annelids are suited for electrophysiological [15–18] as well as behavioural [19–22] studies, and their nervous system is anatomically and histologically well described [23–31]. Potential chemoreceptor proteins have been identified in the first published annelid genomes, which contain homologs for insect receptors (41 Ionotropic Receptor (IRs) and 12 Gustatory Receptor-like receptors (GRs)), but apparently not for mammalian ones (Olfactory Receptors (ORs)) [32, 33]. Despite these advantages, the physiology of chemical sensing in annelids is scarcely known.

Unlike terrestrial annelids such as leeches and earthworms, marine annelids possess elaborate head sensory organs with diverse morphologies [34]. Nuchal organs, paired ciliated cavities located at the back of the head, are considered an annelid synapomorphy [35], and are generally regarded as chemosensory. Based on cell morphology, they are the best candidate chemosensory organs, though no physiological support yet substantiates this claim. Palpae, or palps, the most important head appendages for phylogenetic systematization, have been proposed to be chemosensory based on cell ultrastructure and activity-dependent cell labelling [31,36,37]. Similar claims relying on ultrastructure were made for antennae and tentacular cirri, the two other major types of head appendages [31, 38]. Gross in 1921 has shown that the removal of palps or antennae, and to a lesser extent of tentacular cirri, lengthens the reaction time of the nereidid *Nereis virens* to ionic solutions [19]. Yet, physiological evidence is scarce, and at present no direct experimental proof of chemosensitivity exists for any of these four head organs.

We report here a comprehensive study of head chemosensory organ physiology in the marine annelid *Platynereis dumerilii* (**Figure 1A,B**), which can be easily kept in the lab and is amenable to molecular studies. This species belongs to nereidids, the family regarded as best representing the annelid nervous system ([39], p. 735). Rather than adults, we chose to study early juveniles around the 6-days-post-fertilisation (6 dpf) stage, which are already equipped with nuchal organs, palps, antennae, one pair of tentacular cirri (**Figure 1C**) and display various behaviours. Apart from genetics, they present experimental advantages comparable to those of the nematode *Caenorhabditis elegans,* being transparent, developmentally synchronous, easily obtainable in high numbers, and suitable for whole-body light and electron microscopy. A whole-body atlas of gene expression available for this developmental stage [40] constitutes a unique resource which facilitates the characterisation of cell types. Moreover, a connectomic resource exists at a larval stage that has proved powerful in the reconstruction of whole-body neuronal circuits, notably in the context of phototaxis and ciliated locomotion [41–43].

**Figure 1:**
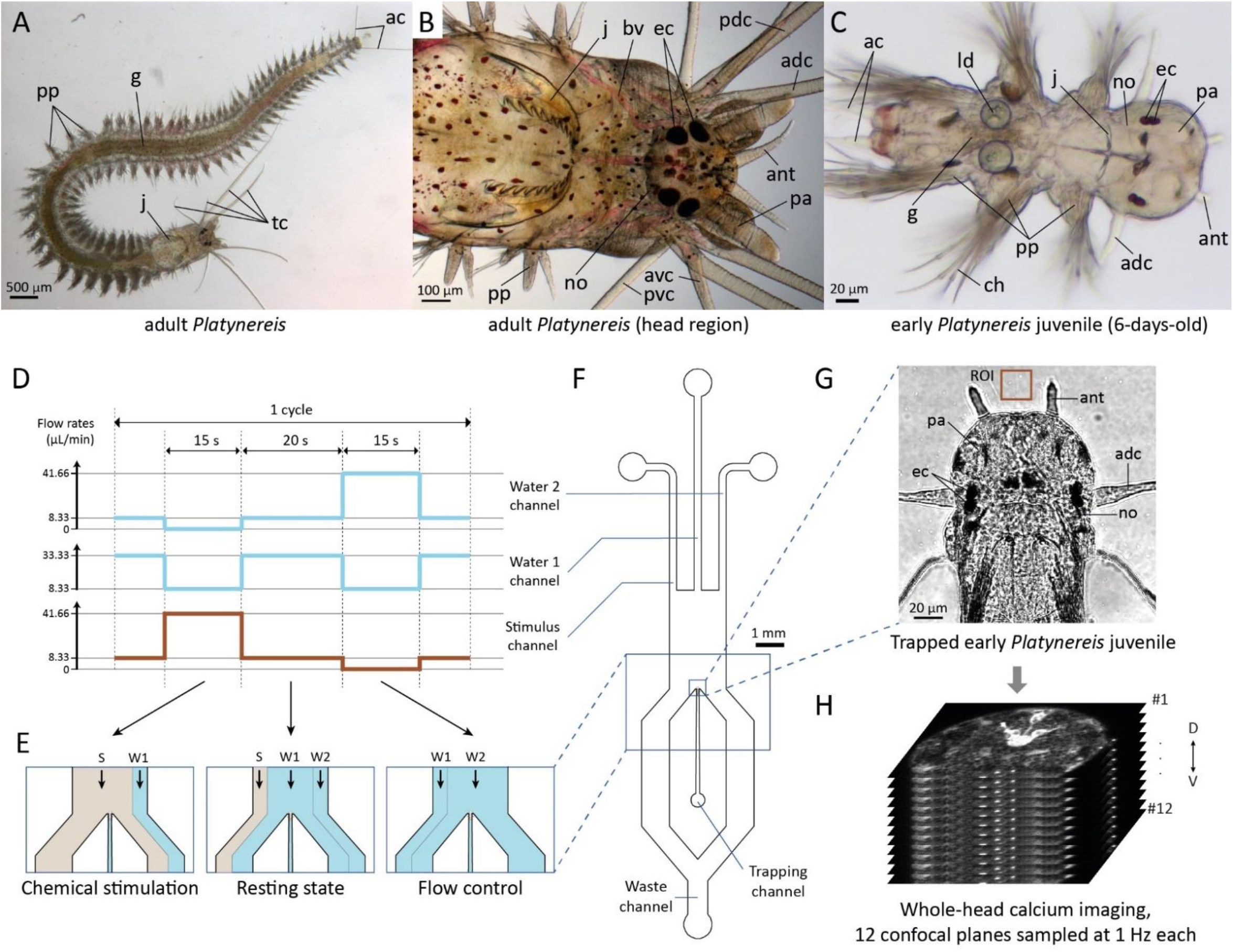
*Platynereis* and the microfluidic device for precise chemical stimulations. **(A-C)** Light microscopy pictures of *Platynereis* at the adult **(A,B)** and early juvenile **(C)** stage, showing antennae (ant), palps (pa) and tentacular cirri (tc, adc, avc, pdc, pvc). Abbreviations: see abbreviations list. A, B, C : © Antje Fischer. **(D)** Channel flow rates used to generate **(E)** the different flow patterns. The sum of the three channels’ flow rates is constant. **(F)** Schematic of the microfluidic device. The three inlet channels are operated by computer-controlled pumps. Single animals are introduced manually in the trapping channel and immobilised at its end. **(G)** An immobilised early juvenile, with its head freely exposed to the sea water flows. Confocal image with transmitted light illumination. **(H)** Calcium signals from trapped animals are recorded with a confocal microscope upon chemical stimulation.

Four stimuli (1-butanol, amyl acetate, glutamate, sucrose) were chosen for their different physico-chemical properties and their likely ecological relevance to *Platynereis*. Short alcohols trigger behavioural reactions in *Nereis* [44]. Amyl acetate can act as a conditioned stimulus for associative learning in the aquatic snail *Lymnea* [45]. Glutamate elicits behavioural responses in *Nereis* [44], and amino acids in general are relevant aquatic chemical cues for various marine animals [12,37,46–49]. Sugars can be degradation products of the polysaccharides contained in plants such as eelgrass and seagrass, or in algae, on which nereidid polychaetes are known to feed [19, 50].

Using a customized microfluidic device for animal immobilisation and precise stimulus delivery, we performed whole-head functional imaging in early juveniles ubiquitously expressing the genetically-encoded calcium sensor GCaMP6s. We found that nuchal organs, palps, antennae and tentacular cirri are chemosensory, though with different degrees of specialisation: for example, antennae responded to all stimulants, while nuchal organs were most sensitive to amyl acetate and sucrose but did not respond to glutamate. We observed a chemically-evoked activity in other regions including the mushroom bodies, which could potentially be involved in learning phenomena. We also described a prominent oscillatory activity in the larval apical organ, however not obviously linked to chemical stimulations. We provide the first direct evidence of chemosensory function in annelid head organs, and lay the ground for future investigations of sensory integration.

## Results

### 1. Establishment of a functional imaging assay system for chemosensation in early *Platynereis* juveniles

We designed a simple microfluidic device, using soft lithography as a fabrication technique and the transparent polymer polydimethylsiloxane (PDMS) as main material. The device is symmetric, has a uniform height of 60 μm and consists of a single chamber in which a constant flow of natural sea water is established. Three inlet channels generate three parallel, non-mixing water streams thanks to a laminar flow regime. Changing the relative flow rates between channels allows to expose the chamber’s centre to any of the streams (**Figure 1D,E**). With one side stream being used to deliver a chemical stimulus and the other to perform a flow control, the device allows to separate the purely chemical sensory input from the mechanical one. An early *Platynereis* juvenile can be immobilised at the end of a central trapping channel (**Figure 1F**), where its head is exposed to the water flows (**Figure 1G; see Figure S1A,B** for a detailed description of the setup).

We performed calibration experiments with a dye to estimate the actual changes of stimulant concentration at the level of the animal’s head (see **Figure S1C and Methods**). With this setup, a stimulus onset (switch from the central stream to a lateral one) can be completed within 0.9±0.3 s, and a stimulus offset (the reverse switch) within 1.0±0.2 s. Stimulus onset and offset are effective with a delay of respectively 3.2±0.5 s and 3.0±0.5 s compared to the pump triggering time.

To survey calcium activity in the entire head, we immobilised juveniles ubiquitously expressing GCaMP6s, and acquired images with a confocal microscope from 12 horizontal, equally-spaced optical sections sampling the whole head volume (**Figure 1H**). With the whole stack being acquired within a second, a 1Hz temporal resolution was obtained for each of the 12 planes, hence the precision of stimulus delivery was deemed satisfactory.

### 2. Ten distinct sets of cells show activity during chemical stimulation experiments

To comprehensively identify head regions active in the context of chemical stimulations, we imaged calcium activity across the entire head in response to four stimulants (see Introduction). For each stimulant, 3 to 4 experiments were conducted for each of 9 animals (only 8 for 1-butanol). A single experiment consisted of three identical cycles: each cycle comprised a chemical stimulus and a flow control, lasting 15 s each and spaced by 20 s resting intervals (Figure 1D).

We observed activity in ten bilaterally symmetrical spots. Four paired region, which we identified as the cell masses of the four presumed chemosensory organs (nuchal organs, palp, antennae, tentacular cirri). We also observed activity in a paired chemosensory region, the lateral region, as well as in the dorsal and ventral mushroom bodies regions. Finally, we also saw activity in three bilaterally symmetric pairs of cells: one in the apical organ area, the eyefront cells and the fronto-dorsal cells. The lateral regions, the eyefront cells and the fronto-dorsal cells, so far undescribed, were named for the present study. Activity was also observed in the nuchal, palpal, antennal and cirral nerves, which were thus included in the analyses. In total, ten distinct sets of cells and four nerves showed observable activity.

To make sense of these signals, we first characterised the tissue context of the responding cells at the 6 dpf stage (**Figure 2, Figure S2**). The **nuchal organs (Figure 2A, Figure S2A)** lie posterior to the eyes and slightly more ventral. Short nuchal nerves and nuchal cavities equipped with ciliated supporting cells can be recognised. In the absence of clear anatomical boundaries, only proximity to the cavity and coactivation with the nerve can allow to attribute a cell to the nuchal cell mass. Most calcium activity was observed in a pair of cells, named ‘revolver cells’ due to their typical shape, but other cells next to the cavity were occasionally active. The **palps (Figure 2B, Figure S2B)** lie ventrally, on each side of the mouth opening, and protrude only very little at this stage. Their mobile tip possesses clusters of cilia. The palpal nerves, short and thick, project to the central neuropil in proximity to the dorsal and ventral roots of the circum-oesophageal connectives. The palpal cell masses, delimited by membrane layers and developing coelomic cavities, constitute the largest cell masses in the head. Active cells were observed rather in proximity to the nerves than to the tips. The **antennae (Figure 2C, Figure S2C)** are slender, frontal organs. Cellular extensions and nervous fibres, but not cell bodies (**Figure S3**), are present inside the appendages, whose surface is equipped with clusters of cilia. The prominent antennal nerves, always identifiable in calcium recordings, project laterally to the central neuropil. The antennal cell masses, anatomically well-delimited, are located at the base of the antennal appendages. Active cells were observed throughout the cell masses. The mobile **tentacular cirri (Figure 2D, Figure S2D**) possess clusters of cilia at their surface and nervous fibres. The cirral nerves project to the circum-oesophageal connectives, not to the central neuropil. The cirral cell masses, future cirral ganglia, are well delimited and occupy a lateral and posterior position in the head. Unlike the antennae, the cirral appendages contain cell bodies (**Figure S3**). Activity was recorded only in cells located at their base.

**Figure 2:**
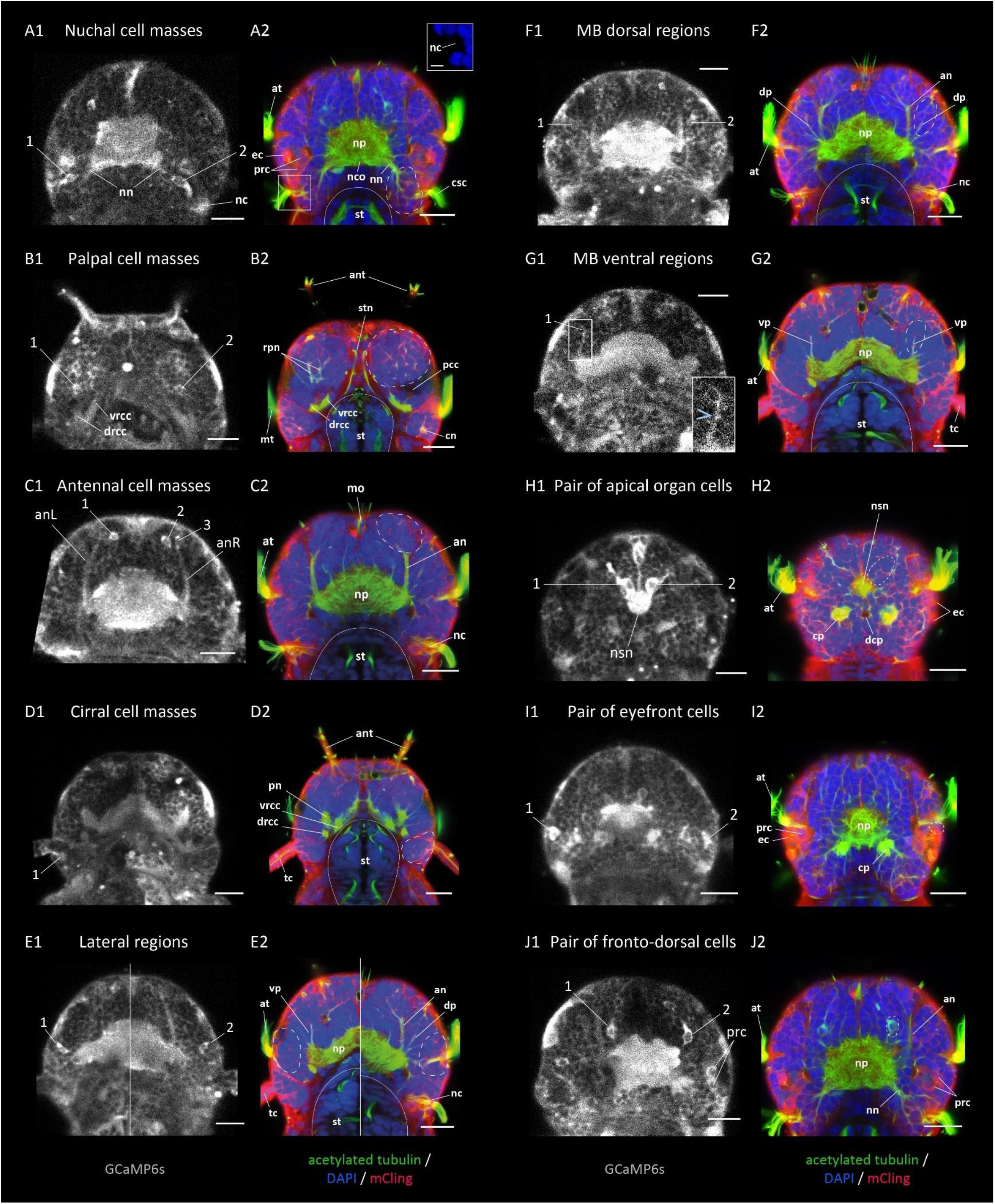
calcium images and anatomical stainings in dorsal view, showing the head regions active in the context of chemical stimulations. **(A1, B1, … J1)** GCaMP6s signal from single confocal planes, averaged over several consecutive time points. Individual active cells are pointed at by numbers. **(A2, B2, … J2)** Acetylated tubulin immunostaining (green), counterstained for nuclear DNA (DAPI, blue) and membranes (mCling, red), averaged over several consecutive confocal planes. The approximate boundaries of the active regions are indicated by white dashed lines, the outlines of the stomodeum by white solid lines. The dorso-ventral position of the planes, as well as lateral views of the anatomical stainings, are shown in Figure S2. Two different horizontal planes have been stitched in E1 and E2. Abbreviations: see abbreviations list. Scale bars: 20 μm for all, 5 μm for inset in **(A2)**.

The **lateral regions (Figure 2E, Figure S2E)** are well delimited, located between the palpal and cirral cell masses, close to the akrotroch. Between one and three active cell bodies were observed in these regions, with no apparent neurite connection to the central neuropil. The **mushroom bodies** (MB) regions consist of probably 15-20 cells each at this stage, organised around two neurite bundles called peduncles: a **dorsal** one, immediately lateral to the antennal nerve (**Figure 2F, Figure S2F**), and a **ventral** one, ventral to the antennal nerve and dorsal to the palpal cell mass (**Figure 2G, Figure S2G**). At 6 dpf, the strong condensation of MB cell nuclei characteristic of adult nereidid brains [51] is not yet apparent, hence precise delimitation of MB cells is not possible. Nevertheless, active cells were observed in immediate proximity to the peduncles, and a coactivation with the peduncle was visible in ventral MB regions (inset in **Figure 2G1**).

Single active cells were observed in the area of the **apical organ** (**Figure 2H, Figure S2H**), an unpaired sensory organ present in annelid and other marine larvae and thought to be involved in their settlement, a crucial life cycle transition [52–54]. These two cells are close to the dorsal head surface, directly anterior to the ciliary band called akrotroch, in proximity to the neurosecretory neuropil known to be associated with the apical organ [55]. They have a flask shape typical for apical organ cells described at 2 dpf [56]. It should be noted that the cilia present dorso-medially at the head surface at 6 dpf do not belong to the apical organ but to another organ of unknown function, the dorsal ciliated pit (see **Figure S2H1**). A second pair of active cells was observed in a position immediately anterior to the eyes, and therefore named **eyefront cells** (**Figure 2I, Figure S2I**). Their activity, though visible in a minority of animals only, was always prominent. A third pair, called **fronto-dorsal cells** (**Figure 2J, Figure S2J**), was observed in a position slightly more dorsal and medial than the antennal nerves, and anterior to the central neuropil. These cells seem to have an axonal projection into the central neuropil, and their shape suggests an anterior cellular extension. A pair of tubulin-rich cells with a similar shape is present at the same position (**Figure 2J2**), but we could not determine whether they are the same.

### 3. Nuchal organs, palps, antennae and tentacular cirri respond differentially to four chemical stimulants

Using four distinct chemical stimulants, we quantified the occurrence of responses for all regions and cells, in each animal, over a long time window following each stimulus onset (**Figure 3A,B**; see Methods). The most obvious responses were those of the antennae, which responded systematically to each of the four stimulants. In contrast, the three other organs showed more differential responses. Nuchal organs were sensitive to amyl acetate and sucrose, and to a lesser degree to 1-butanol, but did not seem to respond at all to glutamate. Palps responded to all compounds, but responses were observed for typically two third of the exposures with glutamate, as opposed to about one third for the other compounds, indicating that palps are particularly responsive to glutamate. In tentacular cirri, responses were seen frequently with glutamate and sucrose, and seldom with 1-butanol and amyl acetate, suggesting that glutamate and sucrose can elicit stronger responses. We performed an analysis of variance to quantify these differences (**Table S1**). A significant difference could be evidenced only for cirral responses to glutamate versus amyl acetate, but more differences would probably become apparent with an increased sample size (here, nine animals per compound). These results show that chemosensitivity in nuchal organs, palps and tentacular cirri is tuned to different types of stimulants.

**Figure 3:**
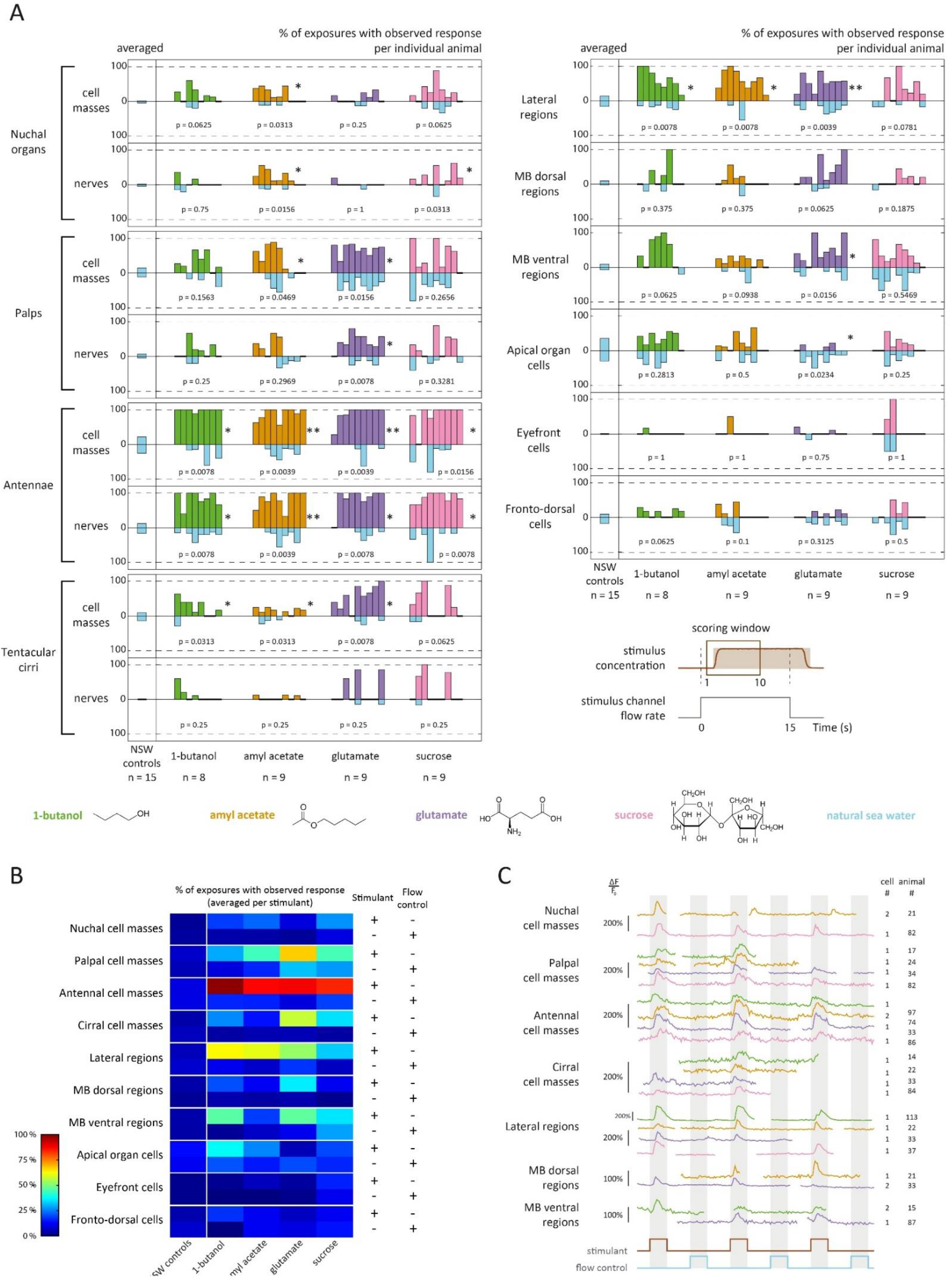
calcium responses of the active regions to the four stimulants. **(A)** Fractions of exposures with observed response for individual animals, shown as mirror barplots for chemical stimulants (green, orange, purple, pink) and flow controls (light blue). For each region, an averaged value obtained in control experiments is shown on the left (separate light blue barplots). p-values and significance levels of a Wilcoxon signed-rank test between responses to chemical stimulants and to flow controls are shown (* = p<0.05, ** = p<0.005). **(B)** Fractions of exposures with observed response, averaged per stimulant. **(C)** Calcium activity traces of individual cells; calcium snapshots and the corresponding regions of interest are shown in Figure S4. Trace colours correspond to the stimulants used.

Different causes seem to account for the non-systematic observation of responses in nuchal organs, palps and tentacular cirri. In nuchal organs, responses were only observed in about 30 % of exposures. The fact that these responses were of high amplitude (typically ΔF/F0 = 100-150 %) and occurred in large cells (diameter of 8-12 μm, as opposed to 5-9 μm in the other regions) excluded that they may have been omitted by our imaging, and suggests that they were conditional. In palps, responses were seen in about 60 % of exposures on average. Since the responding cells were small (<6 μm in diameter) and since abundant muscle fibres and neurites produce calcium signals in this area, we attribute this percentage to technical difficulties in detecting the responses, rather than true biological variability. Responses in the tentacular cirri, whenever observed, occurred in a high fraction of exposures, but were nearly absent in some animals. Since they were overall the weakest responses (typically ΔF/F0 < 50 %), we concluded that the cirri did respond to all stimulants, but the low amplitude of calcium signals did not always allow their detection.

In all organs, the responses were more frequent for the stimulants than for the flow controls (see statistical significance in **Figure 3A**), which confirmed that the observed activity was chemically-evoked. Control experiments without chemical stimulants showed that all organs had comparable levels of responses to natural sea water stimulations as here (**Figure 3A; see Methods**), confirming that responses to flow controls in the present experiments did correspond to a non-chemically-evoked activity. An overall increased activity of the palpal cell masses was nevertheless observed in the particular cases of glutamate and sucrose stimulation.

We next investigated chemically-evoked responses at the single cell level rather than for the entire organs. In palps, antennae and tentacular cirri, we found that most cells which responded to a stimulant did so systematically, and responded only after its onset. Examples are shown for each stimulant in **Figure 3C** (calcium signal snapshots and ROIs are shown in **Figure S4**; more examples of activity traces can be found in [57], ch. IV). These findings indicate that these organs are indeed able to directly detect the onset of such stimulants. In nuchal organs, sucrose was the only stimulant for which we could observe reproducible responses, though only in some of the animals. Two examples of activity traces are shown in **Figure 3C**: one cell responding specifically to the onsets of sucrose, one cell which showed a clear calcium transient after the first onset of amyl acetate and erratic responses upon the second and third exposures. Hence, while the nuchal organs are able to directly respond to the onset of at least some stimulants, such responses do not seem to be robust.

### 4. Chemosensory responses of the mushroom bodies regions and a newly identified lateral region

Similar to nuchal organs, palps, antennae and tentacular cirri, the lateral regions and the mushroom bodies regions showed an enhanced activity directly related to the onset of chemical stimuli (**Figure 3A,B**), which identifies these regions of the early differentiated *Platynereis* brain as part of the chemosensory circuits. Responses were observed in about 65% of exposures for the lateral regions, 40 % for the ventral MB regions, 15 % for the dorsal MB regions. The lateral regions responded to all compounds, though slightly less to sucrose, indicating a broad chemosensitivity as for the antennae. The mushroom bodies regions showed more differential responses, with the ventral ones being more responsive to 1-butanol and glutamate than to amyl acetate and sucrose, and the dorsal ones being mostly responsive to glutamate. Responses to stimulants were more frequent than to flow controls for the three regions, which confirmed their chemically-evoked nature. Only responses of the ventral MB regions upon sucrose stimulation were an exception; in this particular case, another factor than the stimulant may have led to an overall increased activity (three times higher than in control experiments). By inspecting single-cell activity, we could find examples of chemically-evoked responses to all stimulants for the lateral regions, to amyl acetate and glutamate for the dorsal MB regions, and to 1-butanol and glutamate but not sucrose for the ventral MB regions (**Figure 3C**; calcium signal snapshots and ROIs are shown in **Figure S4**). This confirmed that all stimulants were able to elicit responses in the lateral regions, and at least some of them in the two MB regions.

No evidence for stimulant-specific responses could be found in the apical organ cells, the eyefront cells or the fronto-dorsal cells. In the apical organ cells, though we saw responses in about 30 % of exposures to a stimulant, these levels of activity were neither higher than for flow controls nor higher than in control experiments. The same was true for the fronto-dorsal cells though at lower levels, in the order of 15 %. In the eyefront cells, neither the low number of responsive animals (3 out of 35) nor the comparison of their responses to stimulants and to flow controls did allow to conclude at a possible chemosensitivity.

### 5. Responses are observed with a delay in the lateral regions and mushroom bodies regions compared to the major chemosensory organs

Following the observation that seven regions were activated by stimulants, we set out to determine when exactly their responses occurred with respect to the window of exposure. The cumulated distributions of all response times were calculated for each region, by pooling experiments involving all four stimulants (**Figure 4**). The response times correspond to the beginning of calcium transients (see Methods).

**Figure 4:**
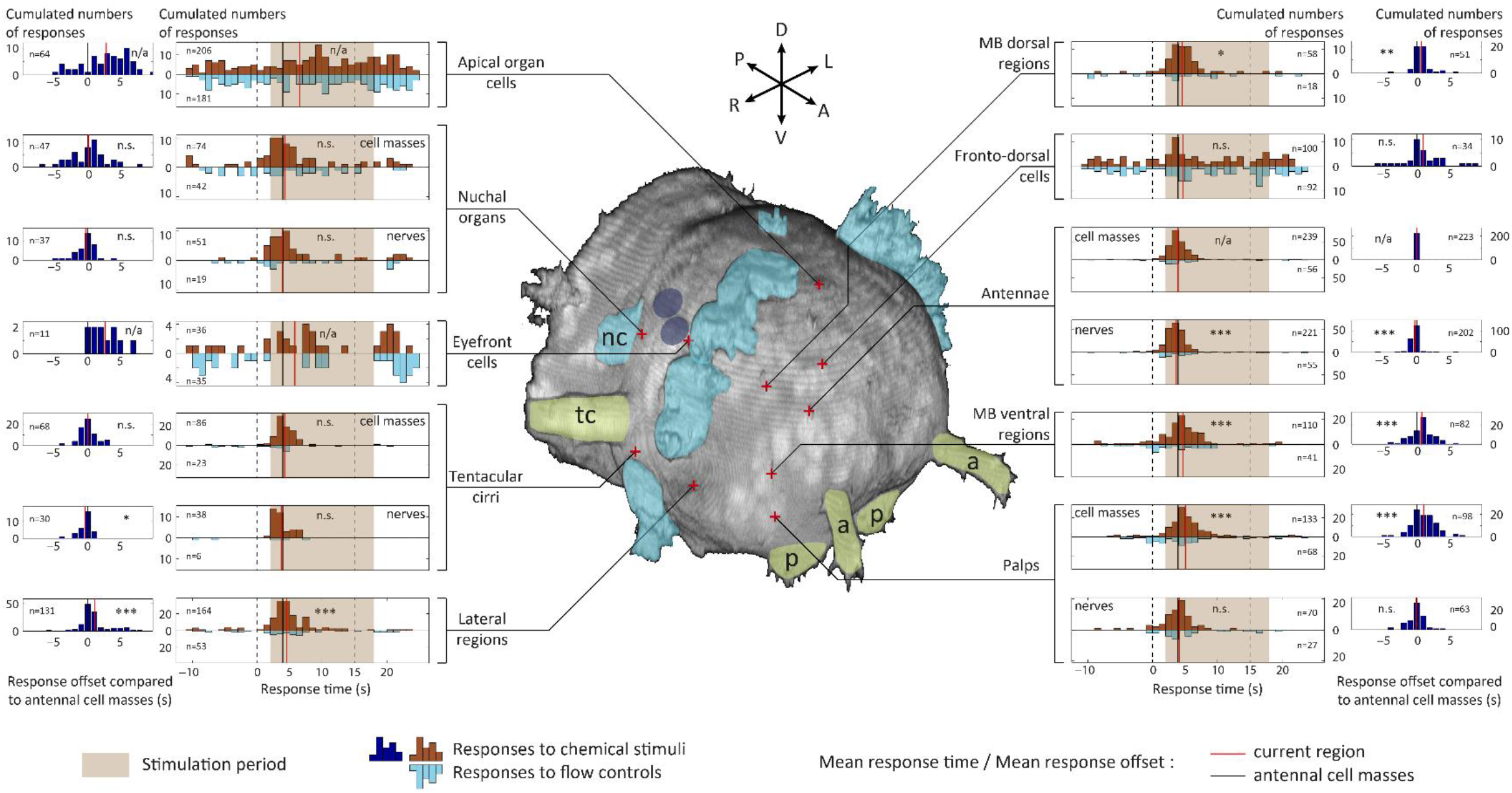
distribution of response times with respect to the stimulation period. Relative response times, with the pump trigger time of the stimulus taken as t = 0 s, are shown in the inner graphs. The total numbers of available responses are indicated. A Student’s t-test was used to compare the mean of the distributions with the mean response time of the antennal cell masses. Delays over the antennal cell masses are shown in the outer graphs. The total numbers of responses for which a delay could be calculated (responses which co-occurred with an antennal response) are indicated. A Student’s t-test was used to compare the mean delay with 0 (n.s. = non-significant, * = p<0.05, ** = p<0.005, *** = p<0.0005, n/a = not applied). The 3D head reconstruction is obtained from the same anatomical stainings as shown in Figure 2. Eyes (dark blue ovals), ciliated structures (cyan) and head appendages (yellow) are highlighted; (a) antenna, (p) palp, (tc) tentacular cirrus, (nc) nuchal cilia.

For the nuchal organs, palps, antennae and cirri, the vast majority of responses took place within 5 s following stimulus onset. The observed variability in these response times was at least partly attributable to variable stimulus onsets as compared to the pump triggering times (**Figure S1C**). Activity outside of the stimulation period was seen more often in the nuchal organs than in the three other chemosensory organs, in agreement with the more erratic responses observed in single-cell activity traces (**Figure 3C**).

For the lateral regions and mushroom bodies regions, responses were likewise observed predominantly following stimulus onset, though the mean response times were slightly delayed. Taking as a reference the responses of the antennal cell masses - the most robust ones, with 239 responses observed overall - we saw indeed a statistically significant delay in the order of 1 s for the mean response time of the lateral regions and of 0.75 s for the two MB regions, but no significant delay or lead for the nuchal organs and cirri (**Figure 4**). In the palps, responses were also observed with some delay for the cell masses, but not for the nerves, indicating that these organs did respond as fast as the antennae.

Complementarily, the offsets of response times with respect to the antennal cell masses were assessed in individual animals, by calculating the actual offset for each response of each region (see Methods). The cumulated distributions of these offsets confirmed the statistical significance of the observed delays (**Figure 4**, outer graphs). A rather wide distribution of offsets was visible for the palpal cell masses and the ventral MB regions, as opposed to the other regions responsive to stimulants. For the lateral regions, the offset distribution seemed to be bimodal, with a main peak at around +1s and a minor one at around +5s (15 % of the responses). A small but significant lead was observed for the antennal nerves over the antennal cell masses (in the order of 0.3 s). The observation that the lateral regions, the two MB regions as well as the palpal cell masses responded with some delay with respect to the overall synchronized responses of the chemosensory organs was robust when offset distributions were alternatively calculated against the cirral cell masses, cirral nerves or palpal nerves (data not shown).

The same analyses applied to the apical organ cells and the eyefront cells confirmed that these cells did not respond specifically to the stimulants, and revealed that their activity was rather uniformly distributed, irrespective of the chemical stimulation period. For the fronto-dorsal cells however, though responses were observed throughout the chemical stimulation period, a peak of responses synchronised to the antennal ones suggested that these cells may show both specific and unspecific responses.

### 6. The apical organ cells show a periodic activity, synchronised with the eyefront cells

Single-cell traces reveal that in 13 out of 35 animals, apical organ cells had slow, large-amplitude calcium fluctuations, as shown in **Figure 5A** (for calcium signal snapshots and ROIs, see **Figure S5**). These fluctuations were synchronous between both cells, as well as with the closely-located neurosecretory neuropil (**Figure 5B**). We found that the eyefront cells, though rarely active, were always synchronised with the apical organ cells (**Figure 5A,B**), which suggests that these two pairs of cells are interconnected. The calcium fluctuations typically followed a period of about 35 s (Animal #29, 30, 33, 37, 40), which corresponds to the periodicity of the alternated stimulations with and without chemical stimulant (see Methods). Although the fluctuations had variable phases compared to the stimulations, this suggests they could have been entrained by the flow stimulations, independently of chemical stimulants; in fact, they can take place in the absence of chemical stimulant (Animal #32). Yet, additional experiments, either without flow patterns or with a different periodicity of flow patterns, would be needed to exclude that these cells had an intrinsic periodic rhythm which accidentally matched the flow period used here. Nevertheless, in two animals, a different pattern of activity was observed, with beginnings of calcium transients correlating with the onsets of chemical stimuli and resulting in similar fluctuations with a period of approximately 70s, i.e. double (Animal #86, 97). This suggests that apical organ cells could still be responsive to chemical stimulants, at least to 1-butanol and sucrose. On the whole, the present experiments do not suffice to conclude whether or not chemical stimulants trigger the oscillating calcium activity in the apical organ cells and eyefront cells.

**Figure 5:**
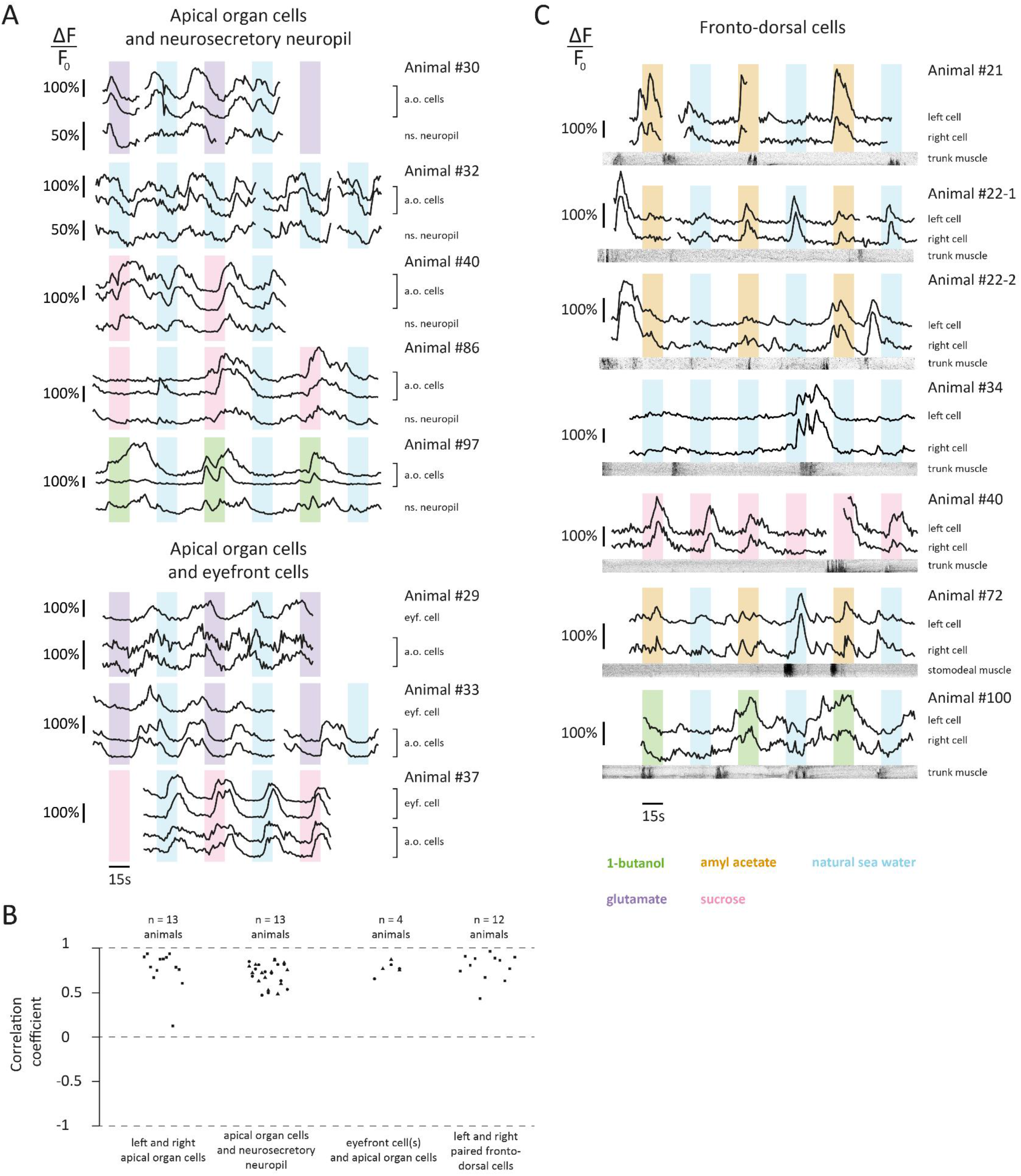
calcium activity of the apical organ cells, eyefront cells and fronto-dorsal cells. **(A)** Individual calcium traces showing synchronisation between apical organ cells and the neurosecretory neuropil (top), and between eyefront cells and apical organ cells (bottom). The stimulations are colour-coded. **(B)** Correlation of the calcium signals between cells. For the neurosecretory neuropil, correlation coefficients are plotted separately for both apical organ cells. **(C)** Individual calcium traces showing responses of the fronto-dorsal cells in relation with stimulations (colour-coded) and with locomotor activity (kymographs). Calcium snapshots and the corresponding regions of interest for **(A)** and **(C)** are shown in Figure S5. Dark parts of the kymographs correspond to movement.

### 7. Activation of the fronto-dorsal cells partially coincides with stimulus onset or termination of locomotor activity

A closer look at single-cell activity confirms that the fronto-dorsal cells do show occasional responses to the onsets of chemical stimuli (**Figure 5C**, Animal #21, 22-1, 40; for calcium signal snapshots and ROIs, see Figure S5), as previously suggested by the distribution of response times. Responses to the onset of the flow controls were also observed (Animal #22-1). Besides, activation of these two cells was seen to follow muscle contraction in the trunk or stomodeum in several animals (Animal #21, 22, 40, 72, 100). The existence of overlapping calcium transients when an episode of movement and a stimulus onset rapidly follow each other (Animal #21, 22-2, 34) suggests that both types of activity may add up.

## Discussion

### *Platynereis* possesses four types of head chemosensory organs

Our calcium imaging experiments based on four types of chemical compounds reveal chemically-evoked responses to these stimulants in nuchal organs, palps, antennae and tentacular cirri (**Figure 3A,B**). Single-cell responses (**Figure 3C**), which probably come from sensory neurons, immediately follow the onset, not the offset, of the chemical stimuli, and are synchronous for the four organs (**Figure 4**). We showed that nuchal organs, palps and tentacular cirri, unlike the antennae, respond differentially to the compounds, which suggests a specialisation of these organs in terms of chemosensory repertoire.

Antennae appear to be central in *Platynereis’* chemosensation, and are probably responsible for the general identification of chemical cues. The fact that their responses were by far more systematic than those of the nuchal organs came as a surprise, since the latter are generally thought to be important for annelid chemosensation, but not the former [24, 35]. The high number of antennal cells already in juveniles suggests that these organs may be capable of chemosensory discrimination. Our attempts to test this further were so far inconclusive. Palps, which are located close to the mouth and were more strongly activated by the amino acid and the sugar (**Figure 3A,B**), may be specialised in the detection of directly food-related chemical cues, as is hypothesised in spionid annelids [22,37,58]. These highly musculated appendages, which adult *Platynereis* use for prehension of food items (personal observations), could also serve in the contact chemoreception of hydrophobic compounds, whose importance is underestimated in marine animals [59, 60]. The tentacular cirri in the present experiments sometimes showed separate response times between left and right side (data not shown). It is likely that in adults, these long organs, which extend in different directions, can provide relevant spatial information about the localisation of chemical cues, at least in relatively still water environments. These tactile organs are also known to be photosensitive and involved in the shadow reflex in *Platynereis* [61]. Hence, their role is probably to collect general multisensory information about objects approaching the head, in order to produce immediate coarse responses. Finally, there is little doubt that the highly conserved nuchal organs play an important sensory role in annelids, hence it is likely that we did not test the most relevant cues for them.

*Platynereis* detects 1-butanol and glutamate, which corroborates behavioural observations made in *Nereis* [19]. Amino acids such as glutamate are general chemical cues in aquatic environments, as is known in fish and crustaceans [62, 63]; other known cues such as nucleotides, steroids and bile acids [64] should be tested in future experiments. The stimulants were presented here at 10 μM, but the presence notably in antennae of responses that slightly precede the earliest possible onset of this concentration (**Figure 4**, see also Figure S1C) suggests that lower concentrations can be detected. The fact that 1-butanol and amyl acetate, which are odorant molecules for humans, act as distance cues in the water for *Platynereis* provides support to the view of Mollo et al. that the traditional categories of ‘olfactory molecules’ and ‘taste molecules’ should be abandoned [60].

Our results in *Platynereis* suggest that marine annelids possess head chemosensory organs with distinct roles, adapted to sets of chemical cues relevant in different situations (feeding, escaping, reproduction, etc.). This is similar to what is known from insect and crustacean chemoreception [7]. Antennae, which seem to be the main chemosensory organs in *Platynereis,* are present in a vast majority of annelid taxa [65] and may thus be of general importance in annelid chemosensation. Though we did not test it here, a distributed chemosensitivity of the body surface is likely for annelids, as suggested by previous behavioural and anatomical studies [19,26,66,67].

### Annelid mushroom bodies as possible chemosensory integration centres

Mushroom bodies, which have long been described in annelid brains [25, 27], have a high anatomical similarity with their homonyms in insects [68]. In fact, similar structures are found in several protostome phyla including flatworms, nemerteans and onychophorans [69], which suggests that they may have been inherited from the last common protostome ancestor’s brain. In insects, mushroom bodies are the place where associative memories are formed, notably with odour stimuli [70–72]. In mammals, this role is endorsed by the pallium [73–75], which includes cortex and hippocampus, and develops from neuroectodermal brain regions expressing similar combinations of transcription factors as in *Platynereis* [76]. We observed cells located in the dorsal and ventral mushroom bodies regions to specifically respond to several chemical stimulants (**Figure 3A,C**). In particular for the ventral regions, coactivation with the mushroom bodies peduncles proved that these responsive cells indeed belonged to the mushroom bodies (**Figure 2G1**, inset). Our observations represent the first physiological data available for annelid mushroom bodies, and firmly establishes them as part of the chemosensory system. Since mushroom body cellular responses were delayed compared to the four major chemosensory organs (**Figure 4**), these responses may indicate the presence of sensory interneurons in the mushroom bodies. This would be in line with a presumed role of mushroom bodies in the representation and integration of chemical cues also in the annelid brain.

### Apical organ cells, eyefront cells and fronto-dorsal cells may be part of the chemosensory circuits

We have identified three distinct pairs of cells in the head whose activity may be partially linked to chemical stimulations. Activity in the apical organ cells did correlate with the onset of chemical stimulations in at least two animals (**Figure 5A**). This preliminary evidence for chemosensitivity calls for further exploration, as apical organs are likely important in the settlement and metamorphosis of marine larvae in general, which they are thought to trigger via the detection of environmental chemical cues [52–54]. The long duration of calcium transients in these cells, as well as their proximity to and coactivation with the neurosecretory neuropil (**Figure 5A,B**), suggest a neurosecretory nature. One can hypothesise that upon detection of appropriate settlement cues, these two cells would adopt periodic patterns of neurosecretory activity, signalling to the animal that settlement can start. Alternatively, calcium fluctuations in these cells may have been merely entrained by the changing flow patterns, as would also be consistent with our data. However, we do not favour this latter hypothesis, as it is hard to explain how this prominent activity would not have been observed in a majority of animals. Whenever the eyefront cells were visible, their activity was tightly synchronised to that of the apical organ cells, hence both pairs of cells may belong to a common chemosensory circuit involved in neurosecretion and/or larval settlement. The pair of fronto-dorsal cells, which are probably neurons due to their morphology and activity patterns (**Figure 2F, Figure S5**), were seen to respond after at least three types of events: chemical stimulations only, chemical stimulations and flow stimulations, locomotor episodes (**Figure 5C**). Hence, their responses to chemical stimulants were not primary sensory responses. We hypothesise that these neurons may have an inhibitory effect, their role being either to prevent a locomotor reaction to external stimuli such as chemical cues, or to stop an ongoing locomotor episode. We hypothesize that these neurons form part of a general circuit for locomotor inhibition receiving inputs from different sensory modalities. As such, they may represent a non-specific part of the chemosensory circuits.

### Variability of responses: biological and technical factors

While antennal responses were strong and systematic for all animals and all compounds, responses in palps and tentacular cirri could not be observed in all animals or for all exposures (**Figure 3A**). Since single-cell responses, though of lower amplitude than in antennae, were always robust (**Figure 3C**), we interpret this fact as a consequence of their response amplitude sometimes falling below our detection threshold. We conclude that these two organs do detect these stimulants, which a more targeted imaging would allow to verify. In contrast, single-cell calcium responses to stimulants in the nuchal organs were hardly repeatable (**Figure 3C**). The typically high amplitude of these responses, whenever observed, rules out an issue of detection threshold. Though we cannot exclude that other compounds may elicit systematic responses, it seems that nuchal organs respond to chemical stimulants in a more conditional manner. However, we were so far unable to tell what influences their responsiveness. For the four chemosensory organs, the nerve was seen to respond with a slight lead over the cell mass (**Figure 4**), even though the former is located anatomically downstream of the latter. We interpret this as an effect of geometry, with local calcium concentrations increasing faster in the axons than in the somata.

### Sensory cell anatomy and physiology is different between nuchal organs and head appendages

Based on our functional imaging data, nuchal organs seem to possess a different physiology than the three other chemosensory organs. In this context, it should be mentioned that nuchal organs are known from electron microscopy studies to possess different types of sensory cells than palps, antennae and tentacular cirri. While cells of the three appendages have sensory cilia that traverse the cuticle and come directly in contact with the environment [23,26,31,38,77–79], sensory cilia of nuchal organ cells sit in a fluid-filled sensory cavity shielded from the environment by specialised cuticular or microvillar layers [80–83]. If these layers affect the diffusion of molecules, changes of chemical composition in the fluid environment may be effective with some delay inside the cavity compared to the surface of appendages. The nuchal organs’ shielded anatomy suggests they may detect the global presence and concentration of ambient chemical cues, but not their rapid concentration dynamics, as the three appendages probably do. A role of these organs in inter-individual communication and the detection of pheromones constitutes a good working hypothesis.

### A microfluidics setup for immobilisation and targeted stimulus delivery

The use of microfluidics for *in vivo* experiments offers several advantages over the immobilisation methods used so far with *Platynereis* juveniles and larvae: gluing, MgCl2 or mecamylamine paralysing, low-melting agarose embedding or slide-coverslip mounting. The present device allows both a reliable animal immobilisation without the use of any chemical agent potentially interfering with the animal’s physiology, and an ecologically-relevant exposure to chemical stimulants. Moreover, stimulant exposures are precise and repeatable, which is key for probing single-cell activity.

Further experiments can be performed with the same setup, notably testing discriminatory abilities of the chemosensory organs between either two concentrations or two stimulants. Any other waterborne stimulus such as pH, salinity, 02 or C02 levels, or even small solid particles, can be used and possibly combined with a precise pharmacological treatment. Identifying a dye undetectable for the animals would allow a more precise monitoring of stimulus timing. Expressing a fluorescent voltage sensor (as in [84]) instead of GCaMP would allow to monitor electrical activity in complement to calcium activity, and whole-head recordings at higher temporal resolutions could be obtained using a light-sheet microscope (as in [85]). The microfluidic device is suited for experiments on 4 to 7 dpf animals, and can easily be adapted for younger stages. Beyond 7 dpf, muscles, hence movement artefacts, are too strong, and either a more elaborate trap or a drug treatment would be needed to achieve immobilisation. The trap design could also be adapted for other marine larvae or small organisms, some of which are suited for microfluidic experiments (personal observations). Though our conclusions would not have been altered, absorption issues documented for PDMS devices may have reduced effective stimulant concentrations [86], hence alternative materials such as COC [87] or sTPE [88] may be preferred to PDMS for future experiments. Microfluidic setups have been successfully used to explore neuronal and motor activity in nematodes and fish larvae [4, 89], our study shows that similar experiments are possible with *Platynereis* juveniles.

### Early *Platynereis* juveniles as a model for the study of annelid chemosensory systems

*Platynereis* sensory organs and nervous systems are representative for annelids, and early juveniles already possess all types of adult chemosensory organs. Our results show that whole-head activity upon precisely controlled chemical stimulations can be imaged in early juveniles. Hence, these juveniles can be used to test the general physiology of annelid chemosensory organs, as monitored by any fluorescent reporter. The availability of a cellular resolution expression atlas [40] in combination with single-cell sequencing and mapping onto the atlas [90, 91] will enable efficient identification of candidate chemoreceptor genes. Candidates would then be validated by combining knock-outs [92] of receptor proteins and functional imaging, and possible *in vitro* deorphanisation of receptors [93]. We have demonstrated that single cells, such as the eyefront cells, fronto-dorsal cells or apical organ cells, can be identified based on their calcium activity patterns (**Figure 2, Figure 3**). Mapping functional imaging data as acquired here onto the gene expression atlas will allow thorough characterisation of such cells. Furthermore, the feasibility of connectomics in *Platynereis* has been proved at the 3 dpf stage [42,43,55,94], and a connectomic effort at 6 dpf is ongoing, which could allow to add circuit information to the molecular and functional data presented here. It will be interesting in the future to investigate *Platynereis’* abilities for sensory integration and associative learning, whose neuronal correlates could be directly studied with the present setup. Finding an involvement of annelid mushroom bodies in associative learning would be of particular interest for the comparative neurobiology of learning.

## Conclusion

We have established that nuchal organs, palps, antennae and tentacular cirri are chemosensory organs in *Platynereis,* responding to an alcohol, an ester, an amino acid and a sugar. This conclusion is likely to extend to annelids, for which similar sensory cells have been detected in electron microscopy. Our results show a capability to differentially respond to multiple chemosensory cues, which opens the possibility of complex chemosensory integration. With our findings, we establish 6-days-old *Platynereis* juveniles as an experimental system for the physiology of annelid chemosensation.

## Methods

### Device fabrication

Standard soft-lithography was used to fabricate the mould [95]. The photomask was designed with AutoCAD (2014 free student version, Autodesk Inc.) and printed at a resolution of 25400 dpi by an external company (Selba S.A., Versoix Switzerland); the source file is available as .dwg file in the electronic supplementary material. The trapping channel’s width linearly decreases from 150 μm to 75 μm at its end, which constitutes the trap. The mould with a uniform height of 60 μm was obtained by spin-coating a silicon wafer (4 inches; Siltronix, France) with a negative photoresist (SU-8 2050, MicroChem Corp., Newton MA). Devices were produced by pouring onto a mould and curing at 65°C for a minimum of 4 h a prepolymer mixture of polydimethylsiloxane (PDMS, Sylgard 184 silicone elastomer kit, Dow Corning Corp.) with a 1:9 ratio of curing agent. PDMS blocks were then irreversibly bound to a 0.17mm glass coverslip (#1871, 24 x 50 mm, Carl Roth GmBH, Germany) by a 1 min treatment in a plasma oven (Femto, Diener electronic GmbH & Co. KG, Germany). For a detailed fabrication protocol, see chapter IV and appendix E in [57].

### Experimental setup and procedure

Filtered Natural Sea Water obtained with 0.22 μm sterile filters (Millipore), plastic syringes (Luer Plastipak, BD, USA), metallic needles (Microlance #20, 302200, BD, USA) and polytetrafluoroethylene tubing (PTFE, TW24, inner diameter 0.59 mm, Adtech Polymer Engineering Ltd, UK) were used in all experiments. A 1 mM stock solution was prepared weekly for each chemical stimulant. Working solutions at 10 μM were prepared on the day of the experiment, and syringes loaded with these solutions were placed in the experimental room 1 h before starting. All solutions were kept at 18°C and handled in glassware, since preliminary experiments had revealed that the animals may detect dissolved substances from plastic containers such as Falcon tubes. Water streams were generated in laminar flow regime (7mm/s, Reynolds number ≈ 0.5). Flow rates of 8.33, 33.33 and 41.66 μL/min were used in the channels (**Figure 1D**), with the total flow rate kept constant (50 μL/min) to minimize pressure changes experienced by the animal. The device was operated by push-pull pumps (AL4000-220Z, WPI Germany, GmbH) which were computer-automated via Micro Manager (version 1.4.21, [96]). Image acquisition and stimulus delivery were synchronised with Auto Mouse Click (MurGee.com), a software for automated mouse actions. A customised metallic chip holder was built to hold the fragile device and facilitate its observation under an upright microscope. For details on the pump automation and the pumping programs, see chapter IV and Appendix D in [57].

### Imaging

GCaMP6s fluorescence excited at 488 nm was detected by a hybrid detector (HyD) set to photon-counting mode in a Leica TCS-SP8 confocal microscope, equipped with a 40x (NA 1.1) water-immersion objective (water was preferred to oil for experimental convenience). Transmitted light images were recorded with a classical PMT detector, from the same excitation light as GCaMP. The head region was imaged in 12 horizontal optical sections (pinhole opened at 6.4 Airy Units) sampling the whole volume at 5 μm intervals. To balance potential biases due to increased signal loss with tissue depth, approximately half of the animals were imaged from the dorsal side and the other half from the ventral side, thanks to adequate trapping. Images were acquired by confocal scanning at 8 kHz (resonant mode, phase X correction 1.32, laser powers 6-28 μW, pixel dwelling time 50 ns).

### Calibration experiments

A green dye (Tartrazine, E102) was dissolved in the two side streams to visualise their moving boundaries. Transmitted light intensity from a 633 nm laser illumination was measured at 10 Hz resolution in a square region of interest, constant in size and position, located just upstream of the animal’s head (brown rectangle in **Figure 1G**). Minimal and maximal intensities, which according to the Beer-Lambert law corresponded respectively to absence of stimulant and maximum stimulant concentration, were normalised between 0 and 1. Edge detection allowed to quantify the beginning and ending of stimulation onsets and offsets (Figure S1C). Measurements were made successively with 8 trapped animals (4 experiments per animal).

### Animal preparation and handling

*Platynereis* juveniles were obtained from a permanent culture following Hauenschild & Fischer’s breeding protocol [97], and kept at 18°C with 16h light / 8h dark cycles. Calcium imaging experiments were conducted at 18-20°C, between 142 and 177 hours post-fertilisation (hpf), at various times of the day and night. For each chemical stimulant, animals coming from at least two distinct batches were imaged. The calcium reporter GCaMP6s [98] was transiently and ubiquitously expressed by microinjecting *Platynereis* eggs with mRNA (1.000 ng/μL) between 1 hpf and the first cleavage. Capped and polyA-tailed mRNAs were synthesised with the mMESSAGE mMACHINE T7 Ultra Kit (Life Technologies) from a vector obtained from the Jékely lab (pUC57-T7-RPP2-GCaMP6 described in [42]). After micro-injection, eggs were kept in Filtered Natural Sea Water and culture conditions remained unchanged. The device was washed with Filtered Natural Sea Water for 4 min before every new animal was manually introduced with a syringe. Each animal was allowed to rest for 5 min before the experiments, and successive experiments on one animal were performed at 5 to 10 min intervals. After the experiments, the animal could be recovered without damage for potential further observations, by gently flushing it out of the device.

### Response assessment

Cellular or nerve responses were assessed with the human eye from raw recordings. In the absence of specific genetic markers, the attribution of a responsive cell to a region relied on its position, guided by anatomical landmark recognition based on precise reference immunostainings (**Figure 2, Figure S2**). A given region was considered to respond whenever at least one cell was seen to respond in at least one of the region’s two bilaterally symmetric parts – for a given nerve, whenever at least one of the two bilaterally symmetric nerves was seen to respond. The time of occurrence of a response was defined as the beginning of the corresponding calcium transient, and the threshold of visual detection of such an event corresponded to a signal-to-noise ratio of approximately 3:1. To be noted: a permanent activity of the eye photoreceptor cells was induced by the 488 nm laser illumination. All response scorings were performed twice at 6-months interval by the same person, the resulting data are available as .xlsx file in the electronic supplementary material (**Table S2**).

### Quantification of brain activity

#### Occurrence of responses

(**Figure 3**). A scoring window was defined, starting with the earliest possible onset of the stimulant (1 s after the pump trigger, see **Figure S1C**) and lasting 9 s. Responses of the regions were scored for each stimulus exposure, with a 1 counted if at least one response was seen during the scoring window, and a 0 otherwise. The fractions of exposures with an observed response were calculated for each animal and each region, separately for exposures to stimulants and flow controls (**Figure 3A**). The averages of these fractions for each stimulant are shown as a heatmap (**Figure 3C**). For each set of animals assayed with a given stimulus, e.g. for the 9 animals tested with glutamate, a Wilcoxon signed-rank test (significance level α = 0.05) was performed for each region to determine whether the fraction of responses over exposures statistically differed between the chemical stimulant and its flow controls (**Figure 3A**). To quantify response occurrences in the absence of chemical stimulation, control experiments were run: 15 of the animals assayed with chemical stimulants had been beforehand imaged in 1 experiment with both side channels containing Filtered Natural Sea Water only, prior to any introduction of chemical stimulant in the device. The response occurrences were quantified as described above, and a global fraction of observed responses over number of exposures was calculated for each region by pooling responses of the 15 animals.

#### Distributions of relative response times and response offsets

(**Figure 4**). All responses of all 35 animals were pooled for each region, and the cumulated distributions of their times of occurrence with respect to the stimulation period were calculated. A Student t-test (α = 0.05) was performed to determine whether the mean of each distribution differed from the mean response time of the antennal cell masses (all means were calculated over the 9 s-scoring window defined for **Figure 3**). Additionally, for each individual response that co-occurred with a response of the antennal cell masses within this window, the time offset between the two was calculated. The cumulated distributions of these offsets for each region are shown as barplots on **Figure 4** (external graphs) for responses to the chemical stimuli only, not to the flow controls. A Student t-test (α = 0.05) test was performed to determine whether the means of these offsets differed from 0, i.e. whether these regions responded at a different time than the antennal cell masses.

#### Single-cell activity traces

**(Figure 3C, Figure 5A,C)** Movement artefacts on the raw calcium recordings were first corrected in ImageJ (version 1.50a) using the plugin StackReg [99] with rigid body transformations. Whenever needed, several parts of the recordings were registered individually and the traces obtained from each of them subsequently reassembled. Mean fluorescence intensity was then calculated from Regions Of Interest (ROI) drawn manually in ImageJ. Further data analysis was done in MATLAB (2014a student version, The MathWorks, Inc.). Traces were plotted as ΔF/F0, with F0 calculated as the mean fluorescence value over a 10-seconds time window during a resting state, i.e. outside of stimulation periods.

### Immunohistochemistry

Animals collected at the 6 days stage (precisely 144 hpf) were fixed in 4% PFA and 0.1% Triton X-100. Tubulin structures were marked with a monoclonal mouse antibody against alpha-acetylated tubulin (Cat# T6793, Sigma–Aldrich GmbH, Germany, 1:250 dilution) and an Alexa 488 secondary antibody (Jackson Laboratories, USA, 1:500 dilution). Nuclear DNA was stained with DAPI (1:1000 dilution) and membranes with mCLING–ATTO 647N (Synaptic Systems GmbH, Germany, 1:50 dilution). DAPI, tubulin and mCling fluorescence were excited at 405, 488 and 633 nm respectively, and recorded with a Leica TCS-SP8 confocal microscope equipped with a 63x glycerol-immersion objective. For details see chapter III in [57].

## Abbreviations list

ac: anal cirrus;
adc: anterior dorsal tentacular cirrus;
an: antennal nerve;
ant, a: antenna;
at: akrotroch;
avc: anterior ventral tentacular cirrus;
bv: blood vessel;
cc: circum-oesophageal connectives;
ch: chaetae;
cn: cirral nerve;
cp: ciliary photoreceptor;
csc: cilia of the nuchal organ supporting cells;
dcp: dorsal ciliated pit;
dp: dorsal peduncle of the mushroom bodies;
drcc: dorsal root of the circum-oesophageal connectives;
ec: pigmented eye cup;
g: gut;
gc: glandular cell;
j: jaw;
ld: lipid droplet;
mo: mechanosensory organ;
mt: metatroch;
nc: nuchal cavity;
nco: nuchal commissure;
nn: nuchal nerve;
np: main neuropil;
nsn: neurosecretory neuropil;
pa, p: palp;
pcc: palpal coelomic cavity;
pdc: posterior dorsal tentacular cirrus;
pm: palpal muscle;
pp: parapodia;
prc: photo-receptor cells;
pvc: posterior ventral tentacular cirrus;
rpn: roots of the palpal nerve;
sog: sub-oesophageal ganglion;
st: stomodeum;
stn: stomodeal nerve;
tc, c: tentacular cirrus;
vp: ventral peduncle of the mushroom bodies;
vrcc: ventral root of the circum-oesophageal connectives.

## Acknowledgements

We thank Maria Tosches, Kaia Achim, Paola Bertucci (Arendt Lab), Csaba Verasztó, Luis Alberto Bezares-Calderón, Cristina Piñeiro-Lopez (Jékely Lab) for technical help on molecular biology work, microinjections and calcium imaging. We thank Csaba Verasztó and Robert Prevedel for helpful comments on the manuscript. We further thank Nirupama Ramanathan and the Merten Lab (EMBL) for initial training on microfluidic fabrication, Christian Liebig from the MPI Tübingen Light Microscopy facility and all staff from the EMBL Advanced Light Microscopy Facility for continuous support.

## Authors’ contributions

TC and GJ designed the experiments. TC and JD built the setup. TC performed the experiments and analysed the data. WD performed the immunostainings. TC and DA interpreted the results and wrote the manuscript. All authors commented and approved the final manuscript.

## Data accessibility

The datasets supporting the conclusions of this article have been provided as electronic supplementary material. Raw calcium recordings are available on demand.

## Ethics statement

Not applicable.

## Supplementary material

**Figure S1:**
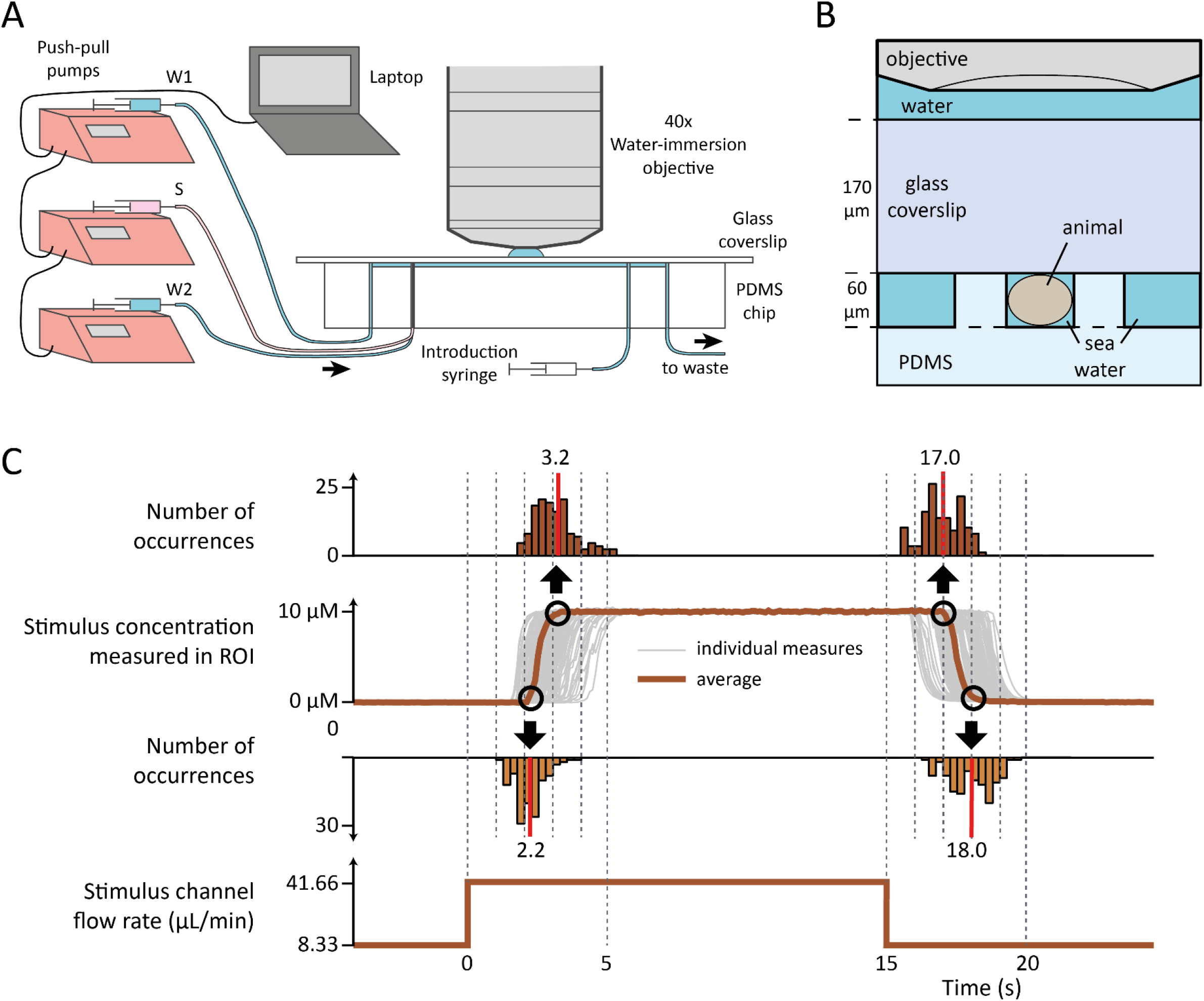
additional descriptions of the experimental setup. **(A)** General schematic showing computer-controlled pumps, tubings and position of the device under the objective. **(B)** Detail of the mounting: cross-section through the microfluidic trap, illustrating the chamber’s height compared to the coverslip thickness. **(C)** Quantification of stimulant concentration over time, as estimated by transmitted light intensity, measured in a square area located immediately upstream of the animal’s head (see Figure 1G). Multiple measurements are shown as grey curves and their average as a thick brown curve. Distributions of the times of occurrence are shown for the four edges of the experimental curves (black circles), which correspond to beginning and end of stimulus onsets and offsets.

**Figure S2:**
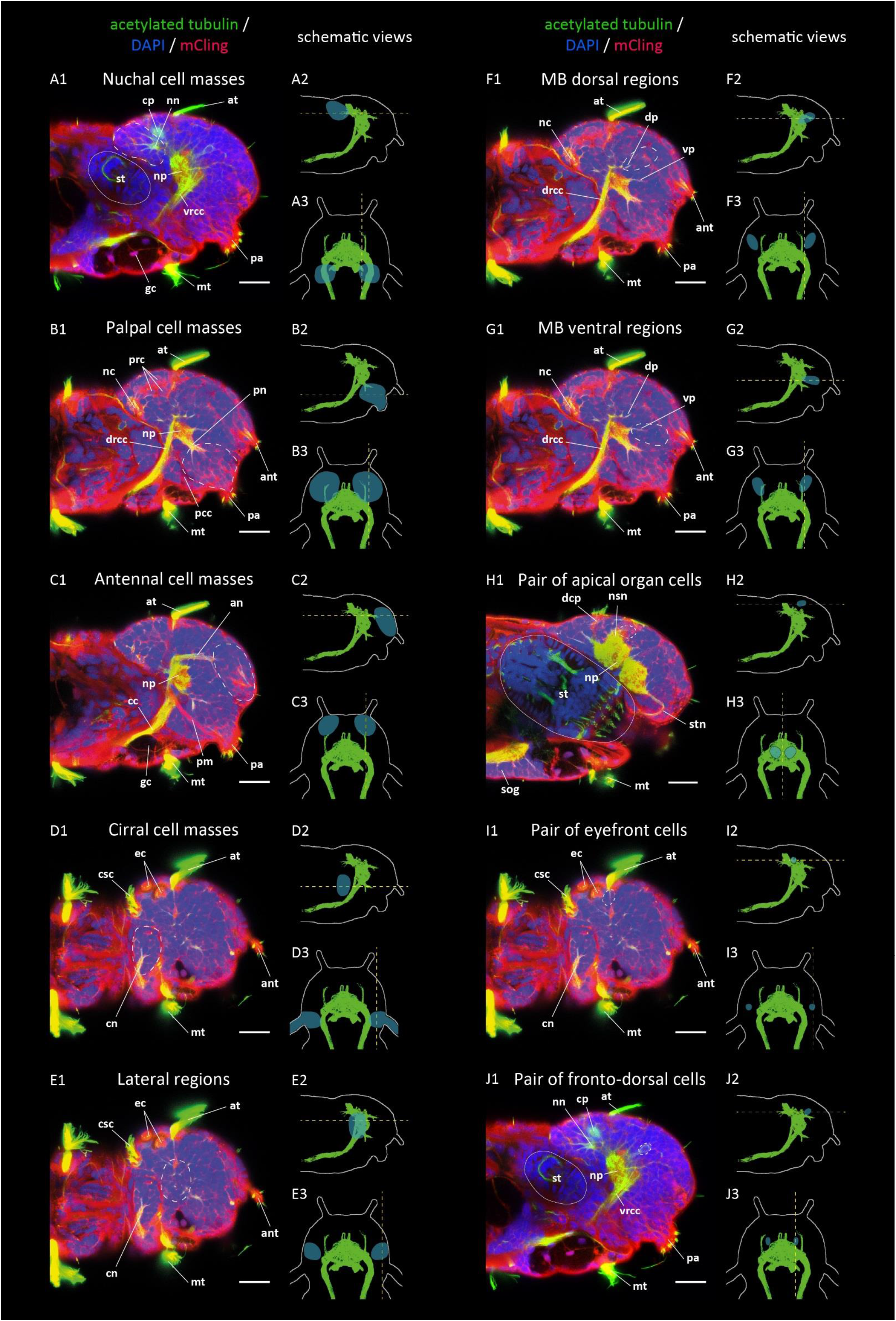
(previous page): anatomical stainings in lateral view, showing the head regions active in the context of chemical stimulations. **(A1, B1, … J1)** Acetylated tubulin immunostaining (green), counterstained for nuclear DNA (DAPI, blue) and membranes (mCling, red), averaged over several consecutive confocal planes. The approximate boundaries of the active regions are indicated by white dashed lines, the outlines of the stomodeum by white solid lines. Abbreviations: see abbreviations list. Scale bars: 20 μm. **(A2, B2, … J2)** Schematic representation of the head in lateral view, showing its outlines (grey), a thresholded maximum projection of the nervous fibres (green) and the area containing the cell bodies of the corresponding region (shaded blue). The position of the horizontal plane shown in the corresponding images in Figure 2 is indicated by a yellow dashed line. **(A3, B3, … J3)** Same in dorsal view. The position of the sagittal plane shown in the adjacent anatomical staining is indicated by a yellow dashed line.

**Figure S3:**
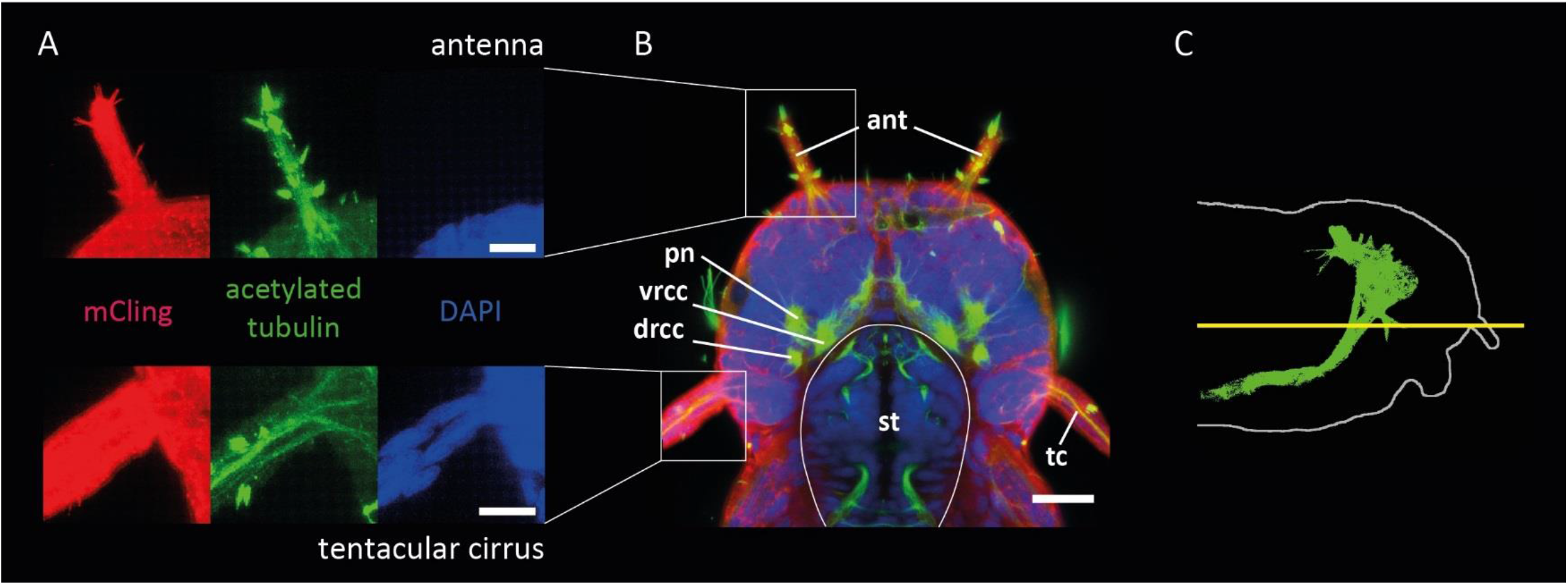
anatomical details of the antennae and tentacular cirri at 6dpf. **(A)** Close-up of head appendages, showing that cell bodies (DAPI nuclear DNA staining, blue) are present in the appendage of the tentacular cirrus (bottom), but not of the antenna (top). Both appendages contain membrane extensions (mCling membrane staining, red), nervous fibres and clusters of sensory cilia (acetylated tubulin immunostaining, green). **(B)** Head overview in a horizontal plane. The outlines of the stomodeum are indicated by white solid lines. Abbreviations: see abbreviations list. **(C)** Schematic representation of the head in lateral view, showing its outline (grey), a maximum projection of the nervous fibres (green). The position of the horizontal plane shown in **(B)** is indicated by a yellow solid line. Scale bars: 10 μm in **(A)**, 20 μm in **(B)**.

**Figure S4:**
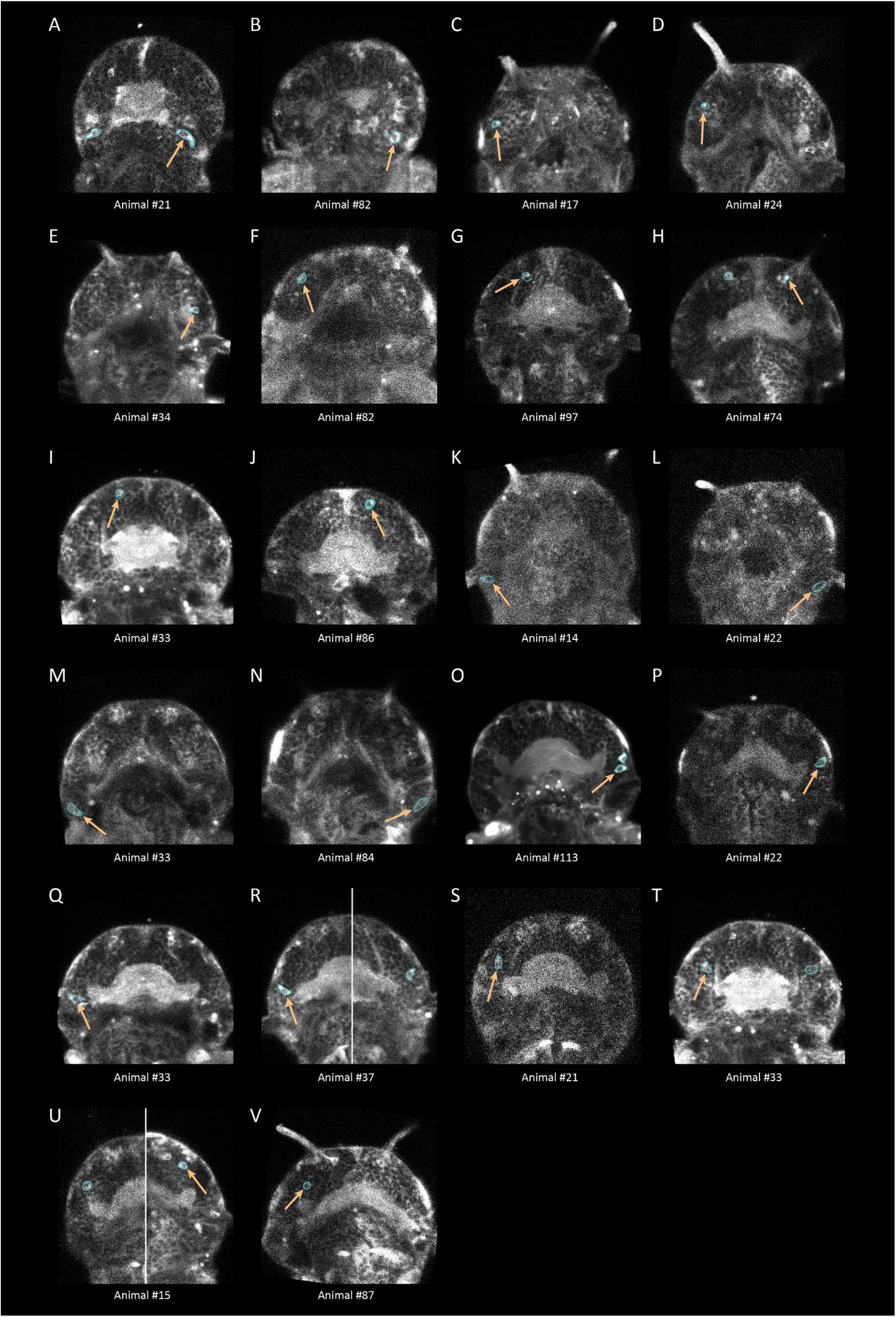
(next page): calcium snapshots and regions of interest used to generate the calcium traces shown in Figure 3C. GCaMP6s signal in single confocal planes, averaged over several consecutive time points. Individual responsive cell bodies are circled in cyan; the orange arrows point at the cells whose trace is shown in Figure 3C. **(A-B)** nuchal cell masses, **(C-F)** palpal cell masses, **(G-J)** antennal cell masses, **(K-N)** cirral cell masses, **(O-R)** lateral regions, **(S-T)** MB dorsal regions, **(U-V)** MB ventral regions.

**Figure S5:**
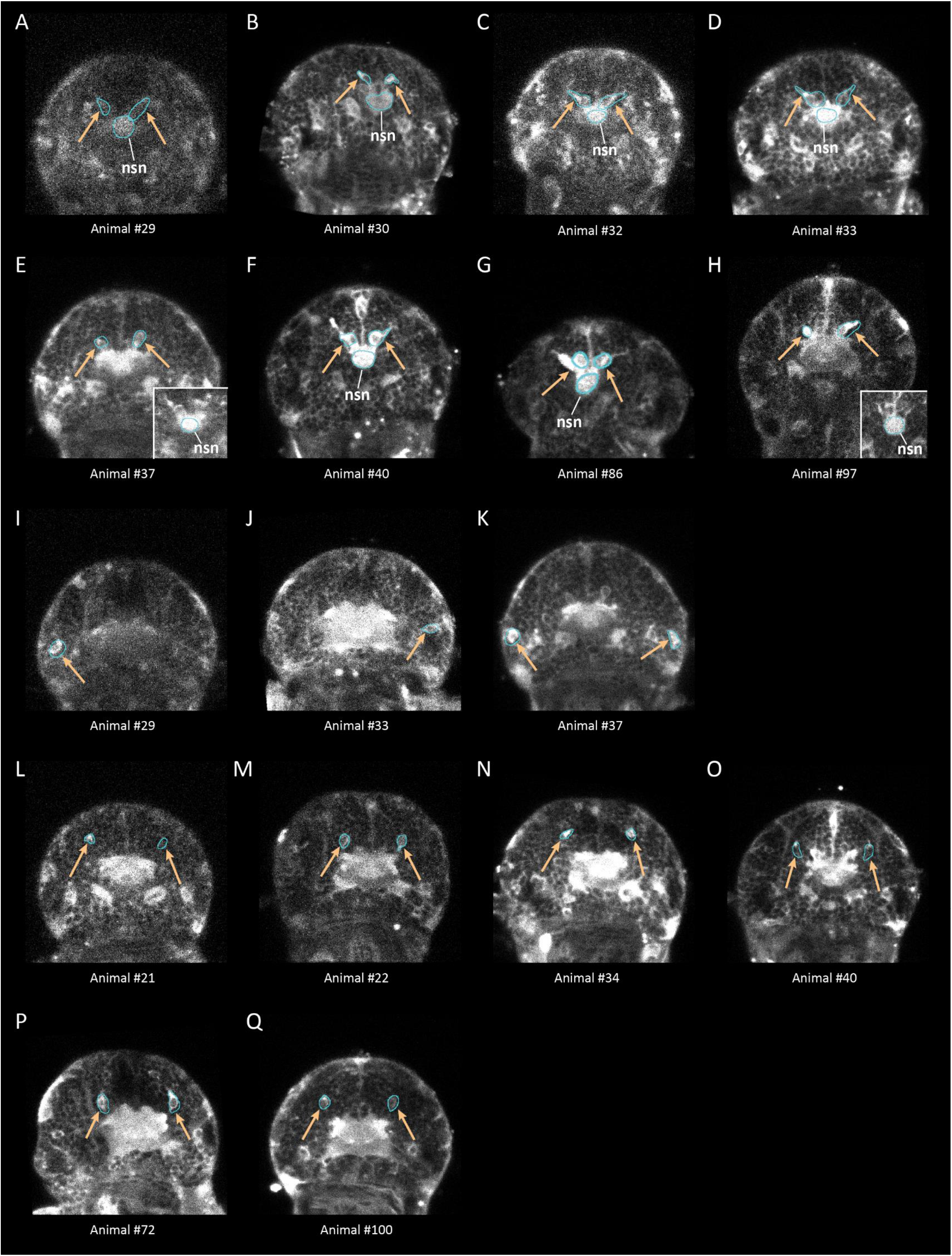
(previous page): calcium snapshots and regions of interest used to generate the calcium traces shown in Figure 5. GCaMP6s signal in single confocal planes, averaged over several consecutive time points. Individual cell bodies or neuropils whose trace are shown in Figure 5A or 5C are indicated by an orange arrow. The insets in **(E)** and **(H)** are taken from a neighbouring confocal plane. **(A-H)** apical organ cells and neurosecretory neuropil, **(I-K)** eyefront cells, **(L-Q)** fronto-dorsal cells.

**Table S1:** multiple comparison test based on one-way ANOVA, for differential responses to stimulants.

|  | groups compared |  | difference between group means | p-value | 95% confidence interval of the true difference of means |  |
| --- | --- | --- | --- | --- | --- | --- |
|  |  |  |  |  | lower limit | upper limit |
| Nuchal organs | 1-butanol | amyl acetate | -1,6488 | 0,9984 | -29,0253 | 25,7277 |
|  | 1-butanol | glutamate | 9,1537 | 0,801 | -18,2228 | 36,5302 |
|  | 1-butanol | sucrose | -8,2625 | 0,845 | -35,6391 | 19,114 |
|  | amyl acetate | glutamate | 10,8025 | 0,6898 | -15,7567 | 37,3616 |
|  | amyl acetate | sucrose | -6,6138 | 0,9054 | -33,1729 | 19,9454 |
|  | glutamate | sucrose | -17,4162 | 0,302 | -43,9754 | 9,1429 |
| Antennae | 1-butanol | amyl acetate | 11,4198 | 0,7255 | -18,2804 | 41,1199 |
|  | 1-butanol | glutamate | 9,9868 | 0,7983 | -19,7133 | 39,6869 |
|  | 1-butanol | sucrose | 14,3519 | 0,5628 | -15,3483 | 44,052 |
|  | amyl acetate | glutamate | -1,433 | 0,9991 | -30,2463 | 27,3804 |
|  | amyl acetate | sucrose | 2,9321 | 0,9925 | -25,8813 | 31,7454 |
|  | glutamate | sucrose | 4,3651 | 0,9761 | -24,4483 | 33,1784 |
| Palps | 1-butanol | amyl acetate | -9,2959 | 0,9321 | -51,5404 | 32,9485 |
|  | 1-butanol | glutamate | -37,788 | 0,0926 | -80,0324 | 4,4564 |
|  | 1-butanol | sucrose | -15,0279 | 0,7698 | -57,2723 | 27,2166 |
|  | amyl acetate | glutamate | -28,4921 | 0,2543 | -69,4752 | 12,4911 |
|  | amyl acetate | sucrose | -5,7319 | 0,981 | -46,715 | 35,2512 |
|  | glutamate | sucrose | 22,7601 | 0,4455 | -18,223 | 63,7433 |
| Tentacular cirri | 1-butanol | amyl acetate | 9,9916 | 0,895 | -28,5666 | 48,5497 |
|  | 1-butanol | glutamate | -31,6707 | 0,1377 | -70,2288 | 6,8875 |
|  | 1-butanol | sucrose | -10,5331 | 0,8796 | -49,0913 | 28,025 |
|  | amyl acetate | glutamate | -41,6623 | 0,0244 | -79,0692 | -4,2554 |
|  | amyl acetate | sucrose | -20,5247 | 0,456 | -57,9316 | 16,8822 |
|  | glutamate | sucrose | 21,1376 | 0,4304 | -16,2693 | 58,5445 |
|  |  |  | p-value > 0.05 |  |  |  |
|  |  |  | p-value < 0.05 |  |  |  |

